# Forest fragmentation, anthropogenic land use, and climate shape sand fly community composition in central Panama

**DOI:** 10.64898/2026.10.26.761809

**Authors:** Juliana Hoyos, Chystrie Rigg, Luis Fernando Chaves, Vanessa Vasquez, Milixa Perea, Jose E. Calzada, Azael Saldaña, Vanessa Pineda, Kadir Gonzalez, Sonia Altizer, Raissa Nogueira de Brito, Adelys Reina, J. P. Schmidt, Susan Tanner, Julie Velásquez Runk, Nicole L. Gottdenker

## Abstract

**Background:** American cutaneous leishmaniasis (ACL) is endemic in central Panama, where rapid land use change is transforming tropical landscapes and potentially altering the ecology of sand fly vectors. Although researchers have widely studied environmental determinants of individual vector species, community-level responses of sand fly assemblages to landscape modification remain poorly understood. We evaluated how landscape composition, fragmentation, climate, and topography influence sand fly community structure in a rural landscape of central Panama.

**Methodology/Principal Findings:** We combined metacommunity analyses with joint species distribution models (JSDMs) to quantify species-specific and community-level responses to climatic variables, land-cover composition, and forest configuration. We collected 1,692 individuals representing 30 species, including nine anthropportunist taxa of epidemiological importance. Sand fly assemblages had strong nestedness, low species turnover, and predominantly positive residual associations. Several anthropportunist sand fly species occurred across heterogeneous landscapes, although their responses to environmental conditions differed among taxa. Land-cover composition, forest fragmentation metrics, and elevation together explained about half of the variance in species occurrence and abundance, with the remainder attributed to climate and spatial, temporal, and community-level random effects. Grassland cover contributed most strongly to sand fly occurrence and abundance, whereas artificial surfaces consistently reduced both metrics. Fragmentation effects varied by species: edge density often increased occurrence and abundance, whereas more forest patches generally suppressed populations. Increased precipitation broadly enhanced occurrence and abundance, whereas higher temperatures negatively affected several taxa.

**Conclusions/significance:** Landscape heterogeneity predictably structures sand fly communities, and different forms of anthropogenic disturbance exert non-uniform effects on vector assemblages. The nested organization of these assemblages and the co-occurrence of several anthropophilic species suggest that multiple potential vector species may be present across heterogeneous landscapes. These findings highlight the need for landscape-informed surveillance and control strategies that prioritize grassland–forest interfaces and other disturbed ecotones where sand fly species of epidemiological importance frequently co-occur.

**Author summary:** American cutaneous leishmaniasis, a vector-borne disease, remains common in Panama and other parts of Latin America. Sand flies transmit the parasites that cause this disease, and deforestation, agriculture, and urban expansion are rapidly changing their habitats. We sampled sand flies in rural communities across central Panama to investigate how climate, land cover, and forest fragmentation shape sand fly communities. Environmental conditions were associated with differences in species composition and abundance. Grasslands and forest edges were associated with higher numbers of some sand fly species, whereas more urbanized areas generally supported fewer sand flies. Sand fly communities showed a nested structure, meaning that species-poor sites tended to contain subsets of the species found in richer sites. Several species of potential epidemiological importance occurred across different landscape conditions, although their responses to environmental change varied. These findings highlight the value of incorporating landscape characteristics into sand fly surveillance and identify disturbed habitats and forest–grassland interfaces as priorities for further ecological and epidemiological investigation. Studies incorporating sand fly infection status, host communities, and human exposure are needed to determine how these ecological patterns relate to *Leishmania* transmission.

## Introduction

Many disease systems are sustained by complex communities of vectors, hosts, and pathogens, not a single vector or pathogen, and landscape structure strongly influences their interactions [1]. Rapid anthropogenic land-use change is reshaping tropical landscapes by altering habitat composition, connectivity, and ecological processes, with important consequences for the emergence and transmission of vector-borne diseases [2]. Consequently, understanding how these landscape changes affect disease systems involving multiple vectors and hosts is a challenge in disease ecology [3].

Tropical landscapes support diverse sand fly communities in which numerous species coexist, and some serve as competent vectors of *Leishmania* spp., the causative agents of leishmaniasis, including American cutaneous leishmaniasis (ACL) [4–5]. Because species differ in ecological niches and vectorial capacity, changes in community composition can alter transmission dynamics and promote the emergence of new disease foci [6–9]. Large-scale deforestation and forest degradation disrupt ecological equilibria, reducing sand fly richness in conserved habitats while promoting the adaptation and proliferation of some sand fly species in peridomestic and disturbed environments [10]. These changes destabilize vector–host interactions, concentrating sand flies near forest remnants and human settlements. For example, in the ‘Deforestation Arc’ in the Brazilian Amazon, researchers found that sand fly-vertebrate host interactions peak when forest cover is moderate, and that vectors and reservoir hosts are most frequently detected where forest and agriculture intermingle. [11]. Urbanization and agricultural expansion may further increase transmission risk by creating suitable breeding habitats and altering ecological conditions that influence vector survival and human–vector contact [12–13]. Accordingly, ACL often occur in fragmented landscapes, where remnant forests embedded within deforested matrices can maintain populations of forest-associated vectors. [14–16].

Panama is a compelling setting for investigating how land use and climate affect ACL vectors because it harbors diverse sand fly fauna, undergoes rapid landscape change, and consistently reports among the highest ACL incidence rates in Central America. Recent data indicate that provinces experiencing intense anthropogenic disturbance (Bocas del Toro, West Panama, Coclé, Panama, and Darién) recorded the highest ACL cases in 2022 [17], highlighting the potential convergent impacts of deforestation, urbanization, and agriculture on *Leishmania* transmission. In Panama, six anthropophilic sand fly species are epidemiologically significant due to their abundance and medical importance: *Nyssomyia trapidoi, Psychodopygus panamensis, Lu. gomezi, Nyssomyia ylephiletor, Lu. sanguinaria,* and *Ps*. *carrerai thula* [18]. While coarse-scale studies, such as Valderrama et al. [19] in Panama, show that some anthropophilic species dominate fragmented forests, whereas others prevail in more conserved habitats, species respond variably to factors like forest connectivity and specific types of anthropogenic disturbance. In contrast, Rigg et al. [20] found that seasonality, particularly the transition to the rainy season, was a stronger driver of local sand fly species diversity than land use or forest cover.

Although environmental effects on sand fly populations are increasingly recognized, most studies have examined individual species or simple land-cover associations rather than community-level dynamics [21]. Recent work has begun to describe how land-use change reshapes sand fly assemblages [22–24], but important gaps remain. Species-focused approaches can overlook co-occurrence structure, trait-related differences in responses, and shared environmental filtering, all of which shape which vector species occur together and may therefore influence human exposure. Few studies have jointly modeled species-specific and community-level responses to landscape composition, forest configuration, and climate while accounting for spatial structure, repeated sampling, and species co-occurrence, and, to our knowledge, none has done so for sand flies in Panama.

Here, we aimed to determine how landscape composition, forest configuration, and climate are associated with sand fly community structure in rural communities of Coclé Province, central Panama. Specifically, we asked: (1) Is sand fly species composition across sites coherent and nested, or does it show species turnover along environmental gradients? (2) How do land-cover composition (grassland, cropland, shrubland, artificial surfaces), forest configuration (edge density and number of forest patches), climate, and elevation relate to the occurrence and abundance of individual species, and do these responses differ between species? (3) After accounting for these covariates, do species co-occur more or less often than expected, indicating shared affinities or unmeasured drivers?

We employed a joint species distribution modeling (JSDM) framework to estimate community- and species-level environmental responses while accounting for spatial structure, hierarchical sampling design, and functional traits such as anthropophily [25]. This approach suits assemblages where multiple vector and non-vector species co-occur across heterogeneous landscapes and may respond idiosyncratically to gradients in forest cover, fragmentation, and human habitat modification. Functional traits included anthropophily, interpreted according to Shaw’s classification [26], which distinguishes anthropportunist species that feed opportunistically on humans among multiple vertebrate hosts and anthroaccidentalist species that rarely bite humans by chance. Throughout the text, we used this classification to highlight the epidemiological importance of some species.

## Methods

### Study Area and Sand Fly Collection

Field collections took place at 72 fixed sampling points distributed across 12 communities (hereafter abbreviated as BLM, CAS, COC, CUT, GUA, MOL, NAN, OAJ, PAL, PAZ, POT, and SJU) located in the province of Coclé, Panama (Fig. 1). Coclé Province, central Panama (08°05′–09°03′ N, 80°02′–80°50′ W), contains mountainous northern regions (200–1600 m a.s.l.) and southern coastal plains, with a humid tropical climate and annual precipitation ranging from 2500–4000 mm. Access to healthcare services is uneven: outside the provincial capital (Penonomé), about a quarter of the population lives within 1 km of a primary healthcare center, half within 3.8 km, while the most remote communities are 46.4 km away [27].

**Fig. 1.**
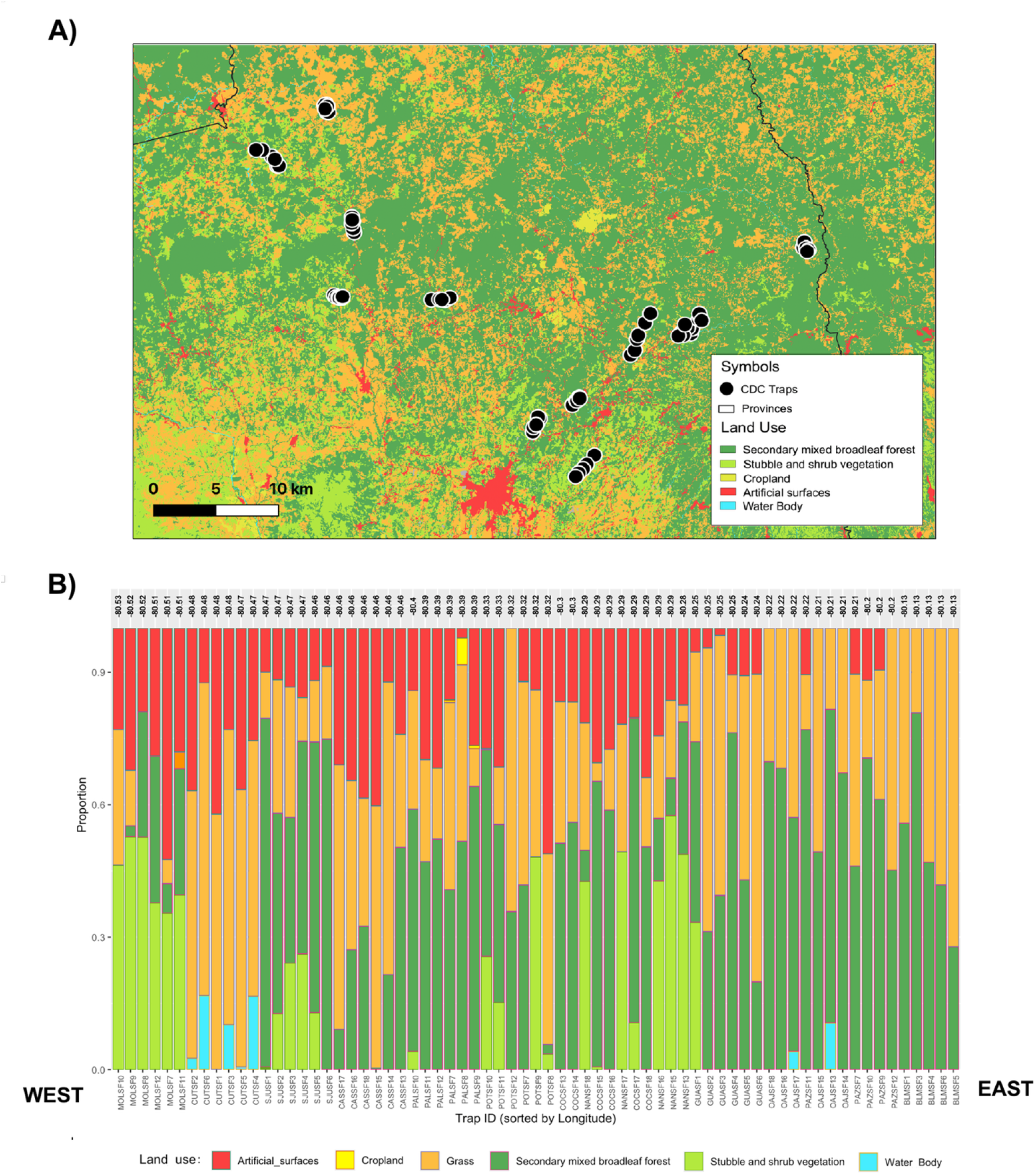
Study area. A) Sand fly surveillance CDC light trap locations, black dots. B) Land cover composition within 300 m of each sampling site across the 72 sampling locations, trap sites arranged from west to east.

Sampling sites were comprised of heterogeneous mosaics of natural and human-modified habitats. Within each community, we collected sand flies using CDC miniature light traps. Traps were installed at standardized collection points and operated during four sampling campaigns: March 2022, November 2022, March 2023, and December 2023. Traps were placed in peridomestic environments. At each sampling event, traps were operated for two consecutive nights at all 72 sampling points (six points per community). We suspended traps 1.5 m above ground level and spaced them more than 200 m apart. Each trap operated for 12 h per night (18:00-06:00). Sampling effort was therefore 8 trap-nights per trap across the four campaigns (2 nights × 4 campaigns), 48 trap-nights per community (6 traps × 8), and 576 trap-nights overall (72 traps × 8), equivalent to 6,912 trap-hours. This gave 288 site-visits (72 traps × 4 campaigns), each representing two pooled trap-nights.

### Specimen Processing and Identification

We euthanized collected specimens with ethyl acetate, preserved them in 95% ethanol, and stored them at −20 °C until processing. Species identification was performed using morphological keys [28–29]. We also supplemented species identification with information compiled by Shaw (2025)[26] to categorize feeding behavior in known species and previous records of *Leishmania* infections. We classified feeding behavior into three categories: anthropportunists, generalist species that feed opportunistically on humans and a wide range of other vertebrate hosts; anthroaccidentalists, host specialists that rarely feed on humans by chance; and zoophilic species, which feed predominantly on non-human vertebrates. This classification was used throughout the study to highlight differences in the epidemiological relevance of sand fly species, recognizing that human exposure depends on feeding behavior and on the environmental conditions influencing human–vector contact.

### Climate and Landscape covariates

Macroclimatic variables were compiled for each sampling site and period using the Giovanni portal of the NASA Goddard Earth Sciences Data and Information Services Center (GES DISC). Precipitation data were obtained from GPM IMERG and temperature from the MERRA-2 reanalysis (NASA’s Global Modeling and Assimilation Office (GMAO)). We assessed temperature conditions using the MERRA-2 reanalysis (M2SDNXSLV; GMAO), from which we extracted daily minimum, maximum, and mean air temperature values. Elevation (m above sea level) was also determined for each trap location.

To characterize the landscapes surrounding sand fly sampling locations, we combined analyses of land-cover composition and forest configuration using remotely sensed land-cover datasets. All analyses were based on the Forest Cover and Land Use 2021 layer developed by MiAMBIENTE [30], based on Sentinel-2 imagery at 10 m spatial resolution (Cobertura Boscosa y Uso de Suelo, Ministerio de Ambiente de Panamá). Geographic coordinates for all CDC light traps were compiled from field records and projected to UTM Zone 17N to align with the land cover raster. Circular buffers of 300 m were generated around each trap, a scale consistent with reported dispersal distances of most Neotropical sand flies [31]. First, within each buffer, we calculated the percentage cover of each land cover class by extracting pixel counts from the raster and standardizing them to proportions. The resulting matrices provided site-specific estimates of land cover composition and percent cover of each of five classes. These data capture variation in 1) secondary mixed forest, 2) shrubland, 3) cropland, 4) grassland, and 5) artificial surfaces, which may influence sand fly populations by affecting host availability, resting sites, and microclimate.

We characterized landscape configuration around each sampling location using the national land cover map. Forest classes were reclassified into binary forest/non-forest categories following MiAMBIENTE and UNCCD classifications (see reclassification table: https://drive.google.com/file/d/1DIhefHNFr2kh-hGKPSdJ234oX5_tiXQo/view). All remaining classes were grouped as “non-forest.” The resulting binary forest/non-forest raster was clipped to the national boundary for subsequent analyses. Within each 300 m buffer, we quantified a suite of forest composition and fragmentation metrics relevant to sand fly ecology and vector–host interactions: 1) forest area (ha), which represents local habitat availability; 2) edge density (m/ha), which capture the extent of forest–non-forest boundaries where vector–host encounters are often concentrated; 3) number of patches (NP), reflecting habitat subdivision; 4) mean patch size (AREA_MN, ha), an indicator of habitat continuity. We delineated forest edges using a Canny edge detection algorithm, and derived patch number and size from an 8-neighbor connected-component analysis. Pairwise correlations among landscape covariates are presented in S1 Fig., and variance inflation factor (VIF) values used to assess multicollinearity are summarized in S1 Table. We excluded covariates with strong correlations or elevated VIF values from the final predictor set to reduce multicollinearity and improve the interpretability of model estimates. Landscape predictors initially considered included the percentages of secondary mixed forest, shrubland, grassland, artificial surfaces, and cropland, as well as edge density and the number of forest patches. After screening for collinearity, the final environmental predictor set included mean air temperature, mean precipitation, elevation, and the selected land-cover composition and forest configuration metrics. We excluded secondary mixed forest because it correlated with grassland, artificial surfaces, and edge density. All continuous predictors were centered and scaled before model fitting.

### Statistical analysis

Sample sufficiency was evaluated with sample-based species accumulation curves, and total richness was estimated using the Chao1 estimator with 1,000 randomizations without replacement to obtain 95% confidence intervals [32]. To standardize comparisons across communities, rarefied richness was calculated by randomly resampling individuals to match the abundance observed in the community with the lowest number of collected specimens. We calculated Pielou’s evenness index (J) to assess the distribution of individuals among species, accounting for seasonal variation. Higher J values indicate more uniform species distributions within communities. Elements of Metacommunity Structure (EMS) were quantified using coherence, turnover, and boundary clumping analyses following [33–34] and implemented with 1,000 null simulations. The procedure yields 1) coherence as the number of embedded absences relative to null expectations (reported with z and p), 2) turnover as observed species replacements along the gradient versus null, and 3) boundary clumping via Morisita’s index with significance testing.

We modeled sand fly communities using joint species distribution models (JSDMs) in the Hmsc package [25]. The JSDM framework lets us partition variation attributable to environmental covariates, spatial structure, and latent species associations, disentangling the roles of habitat configuration and land use. In the context of sand fly-*Leishmania* systems, this ecological lens is critical for anticipating whether anthropogenic environmental change will promote the expansion of anthropportunist vector species and reconfigure community composition in ways that elevate human exposure. We show the complete JSDM framework applied in this study in S2 Fig.

After filtering our initial dataset to retain species present at least 10% of sites, the response matrix comprised 19 species recorded across 72 sites, each sampled during four visits (288 site–visits). This 10% threshold was adopted as a pragmatic criterion for filtering rare species in multivariate community analyses [35], recognizing that species-exclusion thresholds can influence analytical outcomes. We analyzed both abundance data (counts) and presence– absence data (binomial). We modeled species counts with a Poisson distribution (log link), while presences followed a probit distribution. For each site–visit, we compiled two macroclimatic covariates (mean air temperature, mean precipitation) and a suite of landscape predictors computed within 300-m buffers around traps, after applying collinearity screening (VIF analysis and correlogram visualization): percent of shrubland, grassland, artificial surfaces, cropland, and the values for edge density and number of forest patches. All continuous predictors were centered and scaled (mean = 0, SD = 1). We included anthropophily (anthropportunist vs. non-anthropportunist species) as a binary species trait to test whether environmental responses differed by feeding ecology.

Observations were indexed by trap (site), community (12 groups), and visit (four sampling periods). Spatial structure among sites was captured with a spatial random level based on centered–scaled site coordinates (longitude, latitude), with 2–3 spatial latent factors specified a priori. To account for hierarchical grouping and repeated sampling, we included non-spatial random levels for community and visits. Because some traps were located in close proximity, we computed great-circle inter-site distances and defined proximity “blocks” using single-linkage clustering at a 200-m threshold. We added a block-random level to absorb residual correlation among near neighbors not captured by the spatial latent factors. The final study design thus comprised four random levels: site (spatial), community, visit, and “block”. Four MCMC chains were run with 3,000 burn-in iterations and 3,000 posterior samples (thin = 100). Convergence was assessed using Gelman–Rubin diagnostics and trace plots.

Model performance was evaluated separately for the two response variables using metrics appropriate to each observation model: 1) occurrence models based on presence–absence data and 2) abundance models based on count data, as described below.

1. For occurrence models (presence–absence data) analyzed with probit regression, Hmsc implements the Area Under the Curve (AUC), considered a threshold-free classification accuracy [36], and Tjur R^2^ for species-level discrimination [37]. Both metrics measure discrimination, that is, how effectively the predicted occurrence probabilities distinguish between occupied and unoccupied sampling units. For completeness, RMSE was also reported for the occurrence model. RMSE is bounded 0–1 for probit/occurrence models (probability scale), but unbounded (≥0) for Poisson/abundance models (count scale); on link scales it’s still ≥0, expressed in link units. We summarized both species-specific and community-level performances.
2. For abundance models using the Poisson distribution, we assessed model fit using complementary rank-based and error-based metrics. For the abundance model, we evaluated rank-based agreement between observed and predicted counts using the squared Spearman correlation (SR²). To assess model performance conditional on species occurrence, we also calculated C-SR². Whereas SR² evaluates rank agreement across all sampling units, C-SR² assesses how well the model ranks relative abundance among occupied sampling units. Higher values of either metric indicate stronger rank agreement. C-SR² ranges between 0–1, where higher is better. Overall fit across all sites was quantified using root mean square error (RMSE). Conditional predictive accuracy among occupied sites was assessed using conditional root mean square error (C-RMSE), calculated as the square root of the mean squared difference between observed and predicted counts among sampling units where the species was present. Lower C-RMSE values indicate smaller prediction errors and better predictive accuracy for abundance among occupied sites.

In both models, the linear predictor (ηᵢ◻◻) relates the response variable to the explanatory variables and latent effects, determining the expected abundance (λᵢ◻◻) or the probability of occurrence (pᵢ◻◻), respectively. The observation models for abundance and occurrence, together with the corresponding shared linear predictor, were specified as follows:

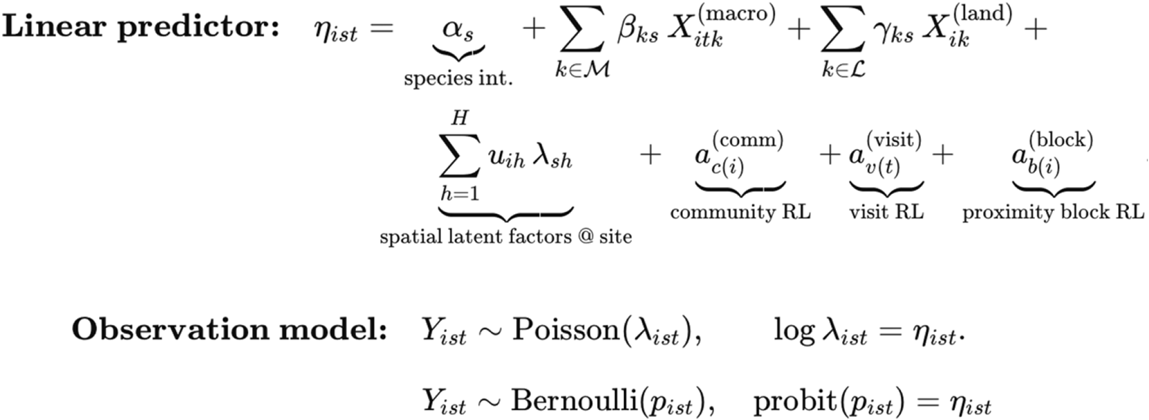

To assess out-of-sample predictive performance while accounting for spatial dependence, we performed grouped K-fold cross-validation with sites as the grouping unit (K = 3). Fold-wise predictions were recomputed from the posterior and evaluated with the same metrics. Site- and community-blocked CVs were also explored in sensitivity analyses. We quantified the relative contribution of predictors and random effects to explained variance on the latent scale using computeVariancePartitioning(). Variance was summarized at both the species level and the community average. S- and predictor-specific effects using posterior means Beta coefficients, and directional support was reported when ≥ 0.90 of the posterior distribution shared the same sign. Residual species associations were estimated from posterior association matrices (Ω) using computeAssociations(). We constructed co-occurrence networks from posterior mean correlations and characterized them using weighted modularity and network nestedness (UNODF).

## Results

### Sand Fly diversity and assemblage structure

We collected 1,692 sand flies representing 30 species across 72 sampling points in 12 communities, where richness ranged from 2–24 species. Some communities showed near-saturation in richness estimates, with small differences between observed and Chao1 richness. In contrast, PAL, POT, and BLM exhibited larger observed-Chao1 gaps, suggesting incomplete sampling and the likely presence of undetected species (S3 Fig.). Richness and diversity increased during wet seasons (S4 Fig.). Zero catches during dry seasons in some communities further underscored strong seasonal effects. Of the 30 species recorded, we classified 9 as Anthropportunist species, many of which are reported to be of high medical importance in Panama (Table 1).

**Table 1.** Abundance and diversity of phlebotomine sand flies collected in Coclé Province, Panama (2022–2023). The table also summarizes reported *Leishmania* infections by sand-fly species and their behavioral categories: Ac = anthroaccidentalist; Ap = anthropportunist; Zo = zoophilic, following Shaw (2025) and Dutari & Loaiza (2014). Blanks indicate unknow behavior.

| Species | Abundance | Feeding preference | Recorded *Leishmania* detection |
| --- | --- | --- | --- |
| *Psychodopygus panamensis* | 502 | Ap | Yes |
| *Lutzomyia gomezi* | 500 | Ap | Yes |
| *Trichopygomyia triramula* | 177 | Ac | Yes |
| *Psathyromyia carpenteri* | 83 | - | No |
| *Nyssomyia trapidoi* | 59 | Ap | Yes |
| *Lutzomyia cruciata* | 53 | Ap | Yes |
| *Nyssomyia ylephiletor* | 48 | Ap | Yes |
| *Lutzomyia aclydifera* | 40 | - | No |
| *Evandromyia saulensis* | 31 | Zo | Yes |
| *Dampfomyia vesicifera* | 31 | - | No |
| *Sciopemyia sordellii* | 24 | Zo | Yes |
| *Dampfomyia vespertilionis* | 18 | - | No |
| *Pressatia camposi* | 17 | - | No |
| *Lutzomyia longipalpis* | 16 | Ap | Yes |
| *Pintomyia serrana* | 14 | - | No |
| *Brumptomyia hamata* | 12 | - | No |
| *Lutzomyia trinidadensis* | 11 | - | No |
| *Lutzomyia lichyi* | 10 | - | No |
| *Lutzomyia sanguinaria* | 9 | Ap | Yes |
| *Pintomyia ovallesi* | 9 | Ap | Yes |
| *Psychodopygus carrerai thula* | 6 | Ap | Yes |
| *Brumptomyia galindoi* | 5 | - | No |
| *Lutzomyia reburra* | 4 | Ac | Yes |
| *Lutzomyia caprina* | 3 | - | No |
| *Psatiromyia shannoni* | 3 | Ac | Yes |
| *Lutzomyia barretoii majuscula* | 2 | Ac | Yes |
| *Lutzomyia guyanensis* | 2 | - | No |
| *Lutzomyia dubitans* | 1 | - | No |
| *Pressatia dysponeta* | 1 | Zo | Yes |
| *Psychodopygus bispinosa* | 1 | - | No |

Sand fly metacommunities had strong positive coherence (z = −9.84, p = 7.5×10⁻²³), reduced turnover (z = −3.69, p = 2.3×10⁻⁴), and pronounced boundary clumping (Morisita’s index = 2.62, p = 6.6×10⁻⁹), consistent with a Nested Clumped Species Loss structure in which species-poor communities represent subsets of richer assemblages. Fig. 2 shows a presence– absence heatmap of sites (rows) and species (columns) ordered by reciprocal averaging (CA1), placing compositionally similar sites and species adjacent to one another and clarifies the primary community gradient.

**Fig. 2.**
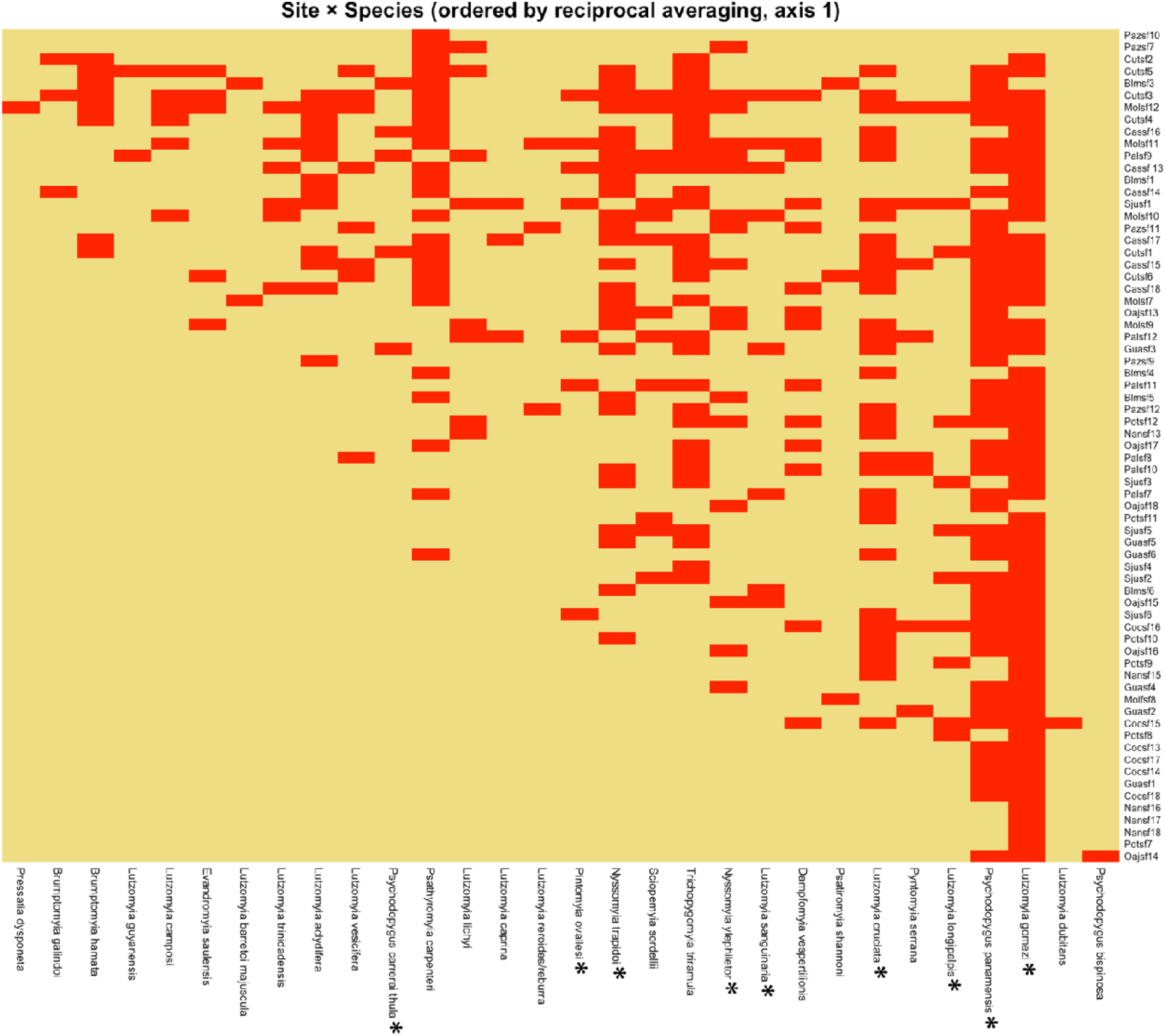
Sand fly distributions (columns) among sites (rows) sampled. A red rectangle within the matrix indicates the presence of that particular sand fly species at the corresponding site. Anthropportunists species are marked with asterisks

### Joint Species model convergence and fit

The joint species distribution models (JSDMs) included 19 sand fly taxa and nine environmental covariates. Secondary mixed forest cover was excluded following collinearity screening because it was correlated with grassland cover, artificial surfaces, and edge density. Consequently, estimated associations with the retained land-cover variables represent conditional relationships with these landscape characteristics, rather than direct contrasts with secondary mixed forest.

Convergence diagnostics indicated that the occurrence model parameters met the predefined criteria (R^ < 1.1 and effective sample size [ESS] > 200; S6 and S8 Figs). In the abundance model, most parameters showed satisfactory mixing, although a subset of species-specific coefficients did not meet one or both criteria (S5 and S7 Figs). We examined these coefficients individually and interpreted abundance estimates associated with parameters showing convergence limitations cautiously.

Occurrence models showed strong in-sample discrimination, with a mean area under the receiver operating characteristic curve (AUC) of 0.916 and a mean Tjur’s (R^2^) of 0.23 across species (Table 2). However, grouped cross-validation indicated lower out-of-sample performance (mean AUC = 0.75, mean Tjur’s (R^2^) = 0.14, and RMSE = 0.259), indicating that predictive performance was more limited than in-sample discrimination. For the abundance model, the mean squared Spearman correlation across all sampling units (SR²) was 0.152, indicating modest agreement in relative abundance rankings. Conditional rank agreement was higher among occupied sampling units (mean C-SR² = 0.417), suggesting the model captured relative abundance differences more effectively where species were present than across all sampling units, including absences. Overall, model performance varied by response type and species (Table 2).

**Table 2.** Model evaluation of JSDMs for sand fly species occurrence and abundance.

| Model Type | Metric | Mean | Median |
| --- | --- | --- | --- |
| Counts | Pseudo R<sup>2</sup> (SR<sup>2</sup>) | 0.152 | 0.100 |
|  | Conditional C -SR<sup>2</sup> | 0.417 | 0.408 |
|  | RMSE (counts) | 0.615 | — |
| Occurrence | Tjur's R<sup>2</sup> | 0.23 | 0.214 |
|  | AUC | 0.916 | 0.928 |
|  | RMSE (probability) | 0.216 | — |

Variance partitioning identified land-cover composition and forest configuration as major contributors to explained variation in both occurrence and abundance (Fig. 3). Grassland cover was the largest individual environmental contributor, accounting for 21% of explained variance in abundance and 14% in occurrence. Artificial surfaces accounted for 7% and 6%, respectively, while cropland contributed 5% and 6%. Climatic variables also contributed to community structure: precipitation accounted for 7% of explained abundance variance and 5% of occurrence variance, whereas temperature accounted for 5% and 7%, respectively. Combined spatial, temporal, and grouping effects (site, visit, block, and community) accounted for approximately 34–39% of explained variance across the two models. These results indicate that landscape composition and configuration were important correlates of sand fly assemblages, while climatic conditions and spatially structured or otherwise unmeasured factors also explained substantial variation.

**Fig. 3.**
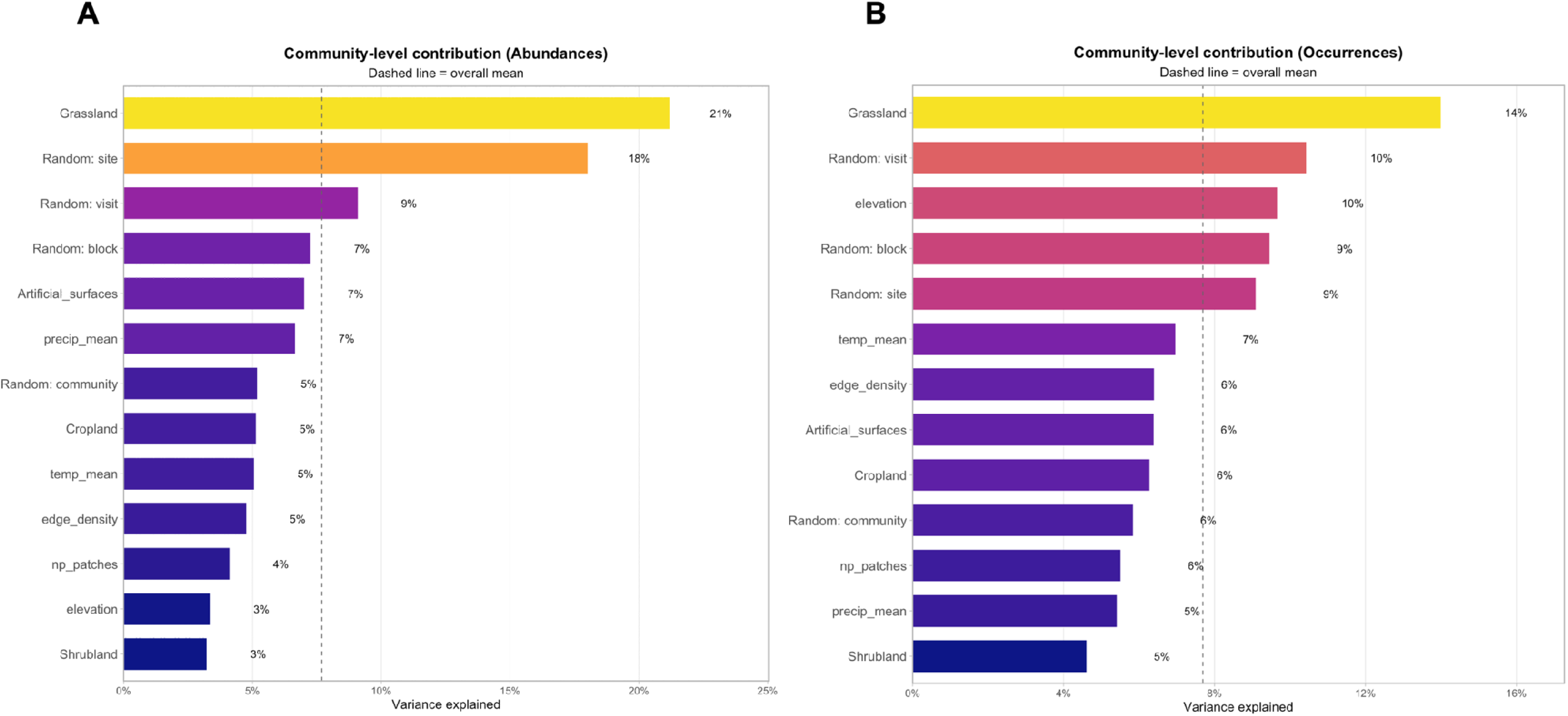
The relative contribution of spatial and environmental variables to the structure of sand fly communities determined from the JSDM for A) Abundance and B) Occurrence data.

### Species-level models fit and responses

Although environmental variables contributed to community-level structure, species responses differed in direction and magnitude (Figs. 4 and 5). Precipitation was positively associated with abundance in *Psychodopygus panamensis*, *Dampfomyia vesicifera*, *Evandromyia saulensis, Pressatia camposi,* and *Pintomyia serrana*. In contrast, higher temperatures were negatively associated with occurrence or abundance in several taxa, including *Da. vespertilionis*, *Ps. panamensis*, and *Ny. ylephiletor*, with posterior support exceeding 90%. Elevation also showed a predominantly negative association with abundance across multiple taxa, indicating that several species tended to be more abundant at lower elevations.

**Fig. 4.**
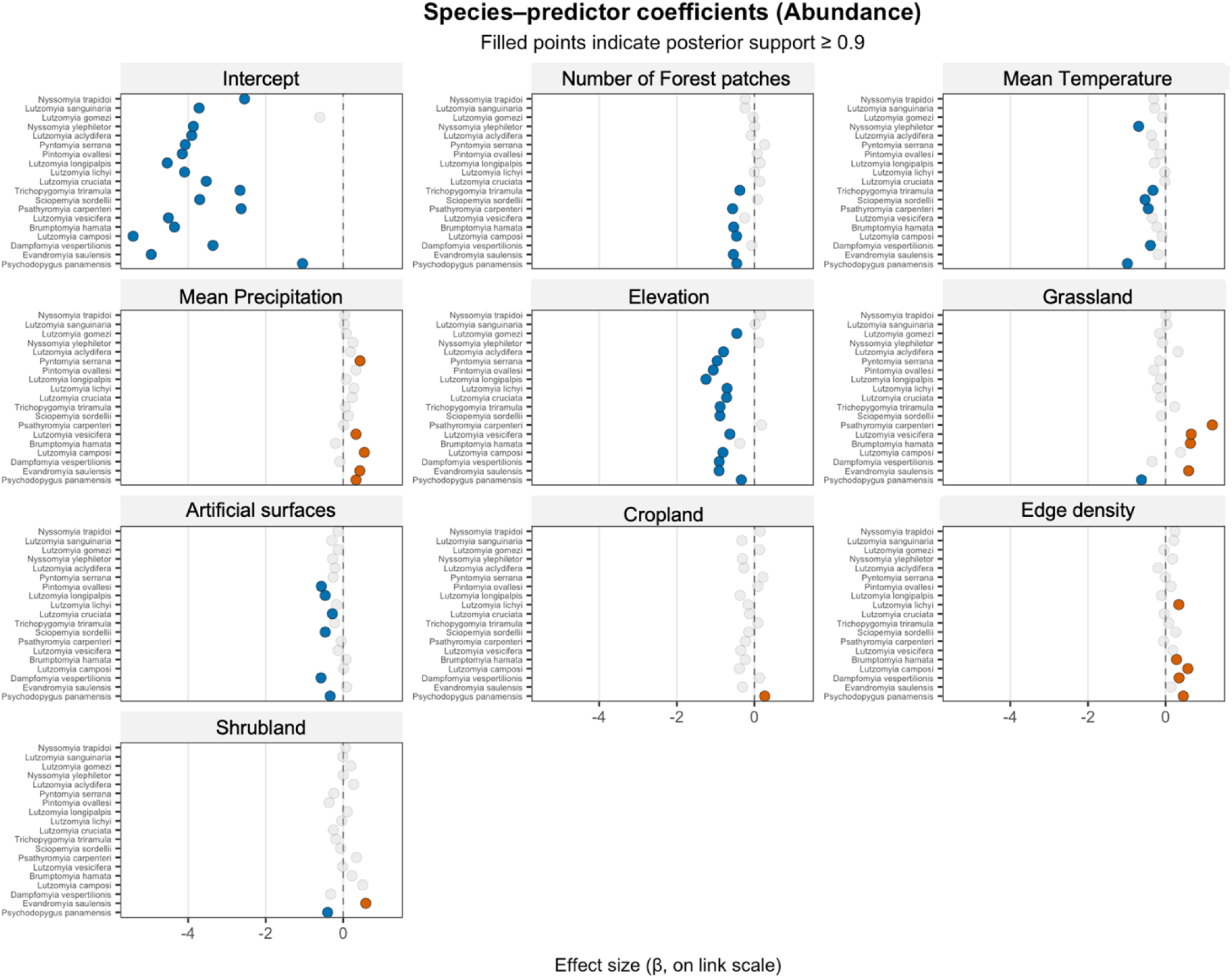
Forest plots summarizing responses, based on β parameters, to all environmental covariates in the JSDM for abundance (CI = 90%). Orange circles indicate positive responses (abundance increases as the values of the environmental covariate increase), while blue circles indicate negative responses (abundance decreases as the value of the environmental covariate increases). White circles indicate CI lower that 90%, thus unclear.

**Fig. 5.**
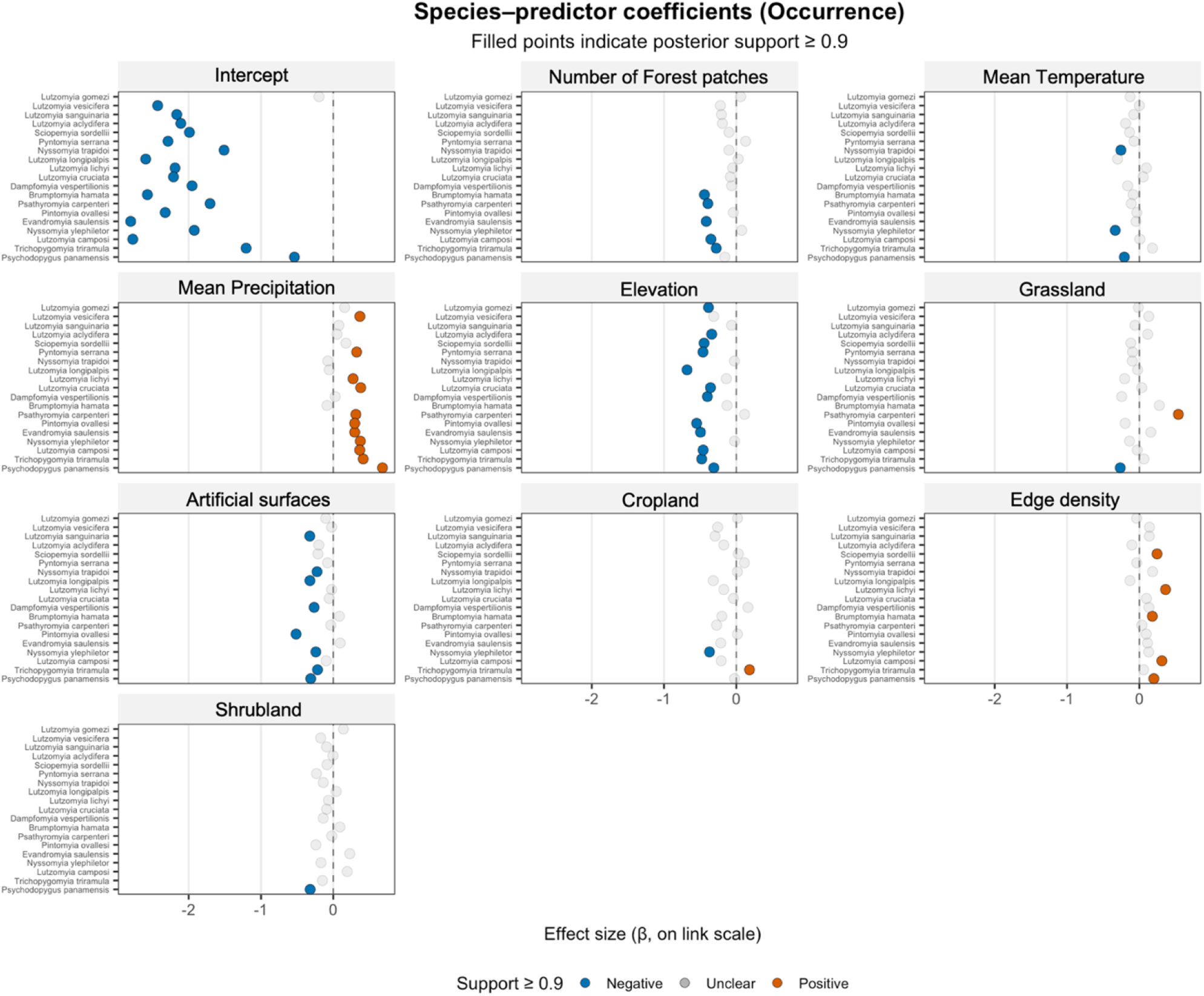
Forest plots summarizing responses based on β parameters to all environmental covariates in the JSDM for occurrence (CI = 90%). Orange circles indicate positive responses (occurrence probabilities increases as the values of the environmental covariate increase), while blue circles indicate negative responses (occurrence probabilities decreases as the value of the environmental covariate increases). White circles indicate CI lower that 90%, thus unclear

Landscape associations were similarly heterogeneous. Edge density was positively associated with occurrence or abundance in several species, including *Br. hamata* and *Da. vespertilionis*. Artificial surfaces were consistently associated with lower occurrence and abundance across the modeled assemblage, whereas a greater number of forest patches was generally associated with lower species counts. Grassland cover produced contrasting responses: it was positively associated with the abundance of *Psathyromyia carpenteri, Dam. vesicifera*, and *Ev. saulensis*, but negatively associated with the abundance of *Ps. panamensis*, with posterior support exceeding 90% for these reported responses. Thus, landscape modification did not elicit a uniform response across sand fly species; instead, different components of landscape composition and configuration corresponded to distinct, species-specific patterns.

Species-level model performance further revealed substantial heterogeneity in how well environmental and random effects accounted for observed patterns (S9–S11 Figs). Conditional rank-based fit for abundance (C-SR²) ranged from 0.015 for *Lu. lichyi* to 0.82 for *Pr. camposi* (median = 0.408). For occurrence, Tjur’s (R^2) ranged from 0.04 for *Lu. sanguinaria* to 0.54 for *Lu. cruciata* (median = 0.214). Marginal AUC values exceeded 0.88 for most species, with *Lu. gomezi* as the principal exception (AUC ≈ 0.74). Together, these results indicate that the models discriminated occurrence relatively well for most taxa, whereas explanatory performance for abundance was more variable and depended strongly on the species considered.

### Spatial species associations and co-occurrence

Residual associations were predominantly positive, suggesting shared ecological affinities among several species. A distinct core of strong residual associations involved the anthropportunist *Ny. trapidoi*, and *Lu. aclydifera*, and *Trichopygomyia triramula* (Ω ≈ 0.81– 0.83). This core was closely associated with *Pa. carpenteri* and *Sc. sordellii* and connected to *Br. hamata* and the anthropportunist *Ps. panamensis* and *Lu. sanguinaria* (Ω ≈ 0.52–0.77).

Approximately 19% of species pairs exhibited moderate positive residual correlations (Ω ≈ 0.35– 0.64). These included *Pr. camposi, Pi. ovallesi, Da. vesicifera, Ev. saulensis*, and the anthropportunist *Ny. ylephiletor*. *Ny. ylephiletor* showed a weak negative residual association with Lu. gomezi (Ω ≈ −0.13), which was otherwise weakly connected to the rest of the network. Most species pairs outside the core exhibited weak or near-zero residual associations, with Pi. serrana and *Lu. cruciata* showing minimal connectivity. *Lu. longipalpis* exhibited the most consistently negative residual associations among the modeled species. However, these associations were weak (Ω ≈ −0.05 to −0.16) and should be interpreted cautiously given the low number of individuals collected (n = 16). Species were grouped according to similarities in their environmental responses and residual associations (Fig. 6).

**Fig. 6.**
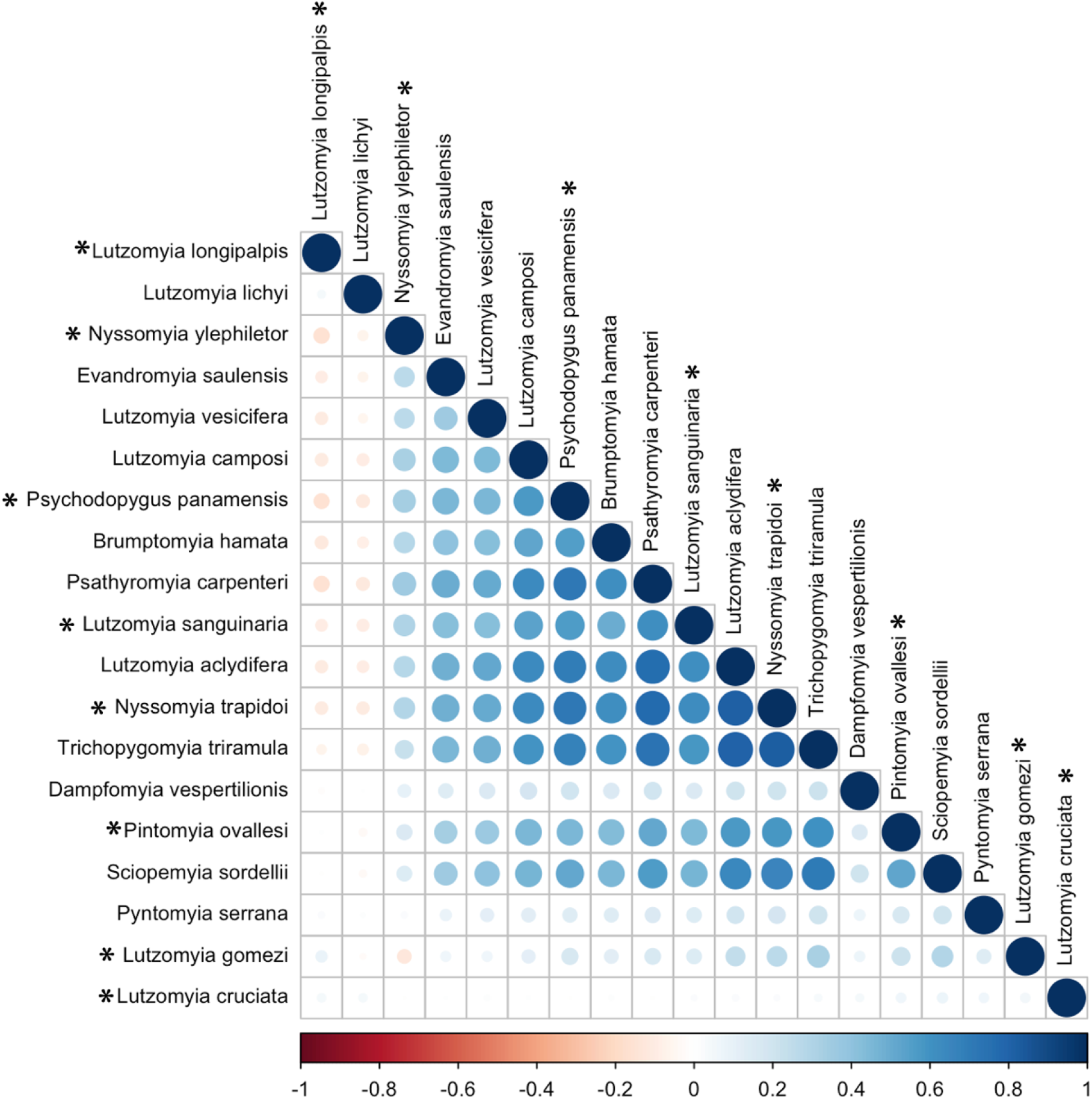
Residual covariance matrix JSDM. Matrix shows mean covariances from the posterior distributions shaded by magnitude and direction, ranging from large negative covariances (deep red) to large positive covariances (blue). Anthropportunists species are marked with asterisks.

At the community scale, high nestedness (UNODF = 80.9; 171 pairs) and negligible modularity (Q=0.015) indicate a core-satellite organization of residual species associations. Species with fewer associations tended to link to subsets of species associated with highly connected taxa, producing a nested network rather than distinct modules. This pattern suggests that community structure is organized around a shared core assemblage, with peripheral species showing diffuse association rather than forming discrete species clusters. Chord diagrams are shown in S12 Fig.

## Discussion

Our results show that landscape composition and configuration strongly shape sand fly community composition in central Panama. We found that land-cover composition and forest fragmentation together explained roughly half of the community signal. Moreover, our approach allowed us to break down the effects of specific anthropogenic activities: Grasslands contributed the largest fixed-effect share for both species abundance and occurrence. Artificial surfaces consistently reduced occurrences, while fragmentation effects (edge density and number of forest patches) influenced species differently, sometimes promoting edge specialists but suppressing others. Climate and topography further modulated assemblages: as precipitation broadly increased occurrence and abundance, higher temperatures reduced them for several taxa. Also, after accounting for environmental factors, positive residual associations suggest that several sand fly species share ecological affinities or respond to environmental and spatial processes not fully captured by the measured covariates. These largely positive associations revealed a tightly connected core of species with high nestedness and weak modularity, reflecting diffuse, nested assemblages rather than sharply partitioned clusters. The predominantly positive residual associations and nested network structure suggest that several sand fly species share environmental affinities or respond to ecological processes not fully captured by the measured covariates. These patterns identify a core assemblage of co-occurring taxa, including species of epidemiological importance. Additionally, species-poor sites formed nested subsets of richer communities, demonstrating high community coherence and low species turnover across the landscape.

JSDMs revealed consistent associations between environmental covariates and sand fly community composition, although predictive performance was lower than explanatory performance. Similar discrepancies have been reported in other insect communities [38–40]. Lower predictive ability likely reflects unmeasured factors, including microhabitat conditions, short-term climatic variability, host communities and biotic interactions, and incomplete sampling at some locations. [38]. The pronounced difference in performance between occurrence and abundance models suggests that stochastic processes, priority effects, or missing environmental covariates may influence sand fly distributions [41]. In addition, the spatial scale at which landscape characteristics are measured may influence their ability to explain community patterns. Our use of local-scale landscape metrics is consistent with previous research showing that landscape characteristics measured within 100–500 m can capture insect responses to forest cover [42]. Nevertheless, because our analyses focused on a specific spatial scale, relationships between landscape characteristics and sand fly communities may differ at finer or broader scales.

The species-specific responses observed here emphasize that landscape disturbance cannot be treated as a single process. Land cover has long been recognized as a major determinant of vector distributions [43][44], yet disentangling the effects of different forms of disturbance remains challenging [45]. Our results show that grasslands, urbanization, and fragmentation affect vector species differently, underscoring the need to evaluate individual landscape attributes rather than broadly categorizing environments as simply “degraded”. In our study, the proportion of grassland within sampling areas was associated with divergent responses, increasing some species’ abundance but reducing others, notably the anthropportunist *Ps. panamensis*. Human management practices such as burning, mowing, grazing, and fertilization strongly influence grassland systems, which often represent transitional stages that precede conversion to croplands. Conversely, some pastures are abandoned and undergo natural ecological succession. Consequently, the effects of grasslands on sand fly communities likely depend on management intensity and successional stage, both of which strongly influence vegetation structure and microclimatic conditions [46]. Grasslands and shrublands may also provide suitable breeding habitats because humidity, organic matter, and livestock feces favor larval development [47]. Furthermore, sand flies are frequently associated with forest–grassland ecotones [48], where livestock, synanthropic reservoirs, and human activities may increase vector-host contact. Similar patterns have been reported in Colombia, where grasslands supported lower diversity but high abundances of a few dominant vector species [49]. The association between grassland cover and the occurrence or abundance of some sand fly species suggests that these habitats may be relevant to vector surveillance. Whether these patterns translate into increased contact among sand flies, reservoir hosts, and humans, or into greater *Leishmania* transmission, remains to be investigated.

The consistent negative association between artificial surfaces and sand fly occurrence and abundance underscores the detrimental effects of urbanization and land conversion on sand fly populations. These patterns likely reflect the loss of suitable breeding and resting habitats that are essential for most sand fly species. Similar reductions in sand flies have been documented in Brazil and other regions where urbanized and heavily modified environments support lower vector densities [50][51]. Less disturbed environments generally provide greater vegetation cover, organic matter accumulation, and favorable microclimatic conditions for sand flies [52]. Rather than eliminating risk entirely, urbanization may shift transmission toward peri-urban interfaces where residual vegetation and domestic animals remain available.

Fragmentation effects can become increasingly important once habitat cover declines below critical thresholds, thereby reducing connectivity and species persistence in species-specific ways [53]. Consistent with this expectation, fragmentation metrics produced contrasting responses among sand fly species in our study, highlighting the importance of considering both landscape composition and configuration. Edge density showed predominantly positive effects on sand fly occurrence and abundance, suggesting that the ecological consequences of fragmentation may be driven strongly by the creation of forest-agricultural interfaces. Such edge habitats alter microclimatic conditions and resource availability, often favoring disturbance-tolerant and generalist taxa [54][55]. The positive responses observed here suggest that these environments provide access to hosts, breeding substrates, resting sites, and favorable microclimates. Similar patterns have been reported in in other insect communities, where ecotones facilitate resource use by both forest and matrix species [42]. Forest edges were associated with increased occurrence or abundance in several sand fly species, suggesting that these interfaces may be useful targets for further ecological and epidemiological investigation.

In contrast, more forest fragments negatively affected several sand fly species, suggesting that excessive habitat discontinuity reduces community diversity. Other insect assemblages show similar patterns, with highly fragmented landscapes supporting fewer species than continuous habitats [56]. Nevertheless, fragmentation effects remain scale-dependent and may differ among diversity components and species, reinforcing the need to evaluate fragmentation at the individual species level rather than relying solely on community-wide responses [53][57].

Our results indicate that sand fly communities in central Panama form a nested metacommunity characterized by clumped species loss, in which species-poor sites represent subsets of richer assemblages. This pattern suggests strong environmental filtering and limited species replacement, with medically important vectors such as *Lu. gomezi*, *Ps. panamensis*, and *Lu. longipalpis* occurring across a broad range of environments, whereas species such as *Ps. carrerai thula* were restricted to the most species-rich sites. Similar patterns have been reported in Venezuela, where anthropportunist sand fly species exhibited broad distributions and minimal turnover among communities [4].

Although higher sand fly diversity is generally associated with conserved and structurally complex habitats, anthropportunist species appear capable of persisting across diverse land-use mosaics. In the Yucatán Peninsula, anthropportunist species remained dominant along gradients of land-use change while more specialized taxa declined [58]. Similarly, Chaves and Añéz [59] reported nested distributions in the Venezuelan Andes and suggested that anthropophilic and zoophilic species exploit overlapping resources. In Panama, *Ps. panamensis* and *Lu. gomezi* frequently co-occur across heterogeneous landscapes [60][20], highlighting the ecological plasticity of major ACL vectors and their ability to occupy both conserved and disturbed habitats. Similar broad spatial distributions have also been observed in other vector communities [61].

Vector metacommunity structure has important implications for pathogen transmission. Unlike *Culicoides* communities, where species replacement along environmental gradients is common [62], sand fly assemblages in Central America appear characterized by low modularity and diffuse associations among medically important species. Whether nested, multispecies assemblages influence the persistence of *Leishmania* transmission cannot be determined from these data and should be tested with infection and host data. Finally, predominantly positive residual associations suggest that several sand fly species share ecological affinities or respond to environmental and spatial processes not fully captured by the measured covariates. The high nestedness and low modularity of the residual association network indicate a relatively connected assemblage structure rather than sharply separated groups of species. Similar patterns have been observed in other insect communities, where apparent segregation largely disappears after controlling for habitat conditions [63][64][65].

### Study limitations

Although the models captured a substantial portion of variation in species occurrence, they explained only modest variation in species abundance, suggesting that stochastic processes, unmeasured environmental variables, and fine-scale habitat features not included here may play significant roles in shaping sand fly community structure. Convergence diagnostics were less satisfactory for the abundance model than for the occurrence model. Although most regression coefficients met the predefined criteria (R-hat < 1.1 and effective sample size [ESS] > 200), a subset did not meet one or both thresholds. Consequently, species-level abundance estimates associated with these parameters should be interpreted cautiously (S5–S8 Figs). Additionally, short-term climate variability and host community dynamics were not directly incorporated, which could further influence vector distributions and human exposure.

## Conclusions

Sand fly assemblages in these rural communities were nested subsets with predominantly positive residual associations, and several anthropophilic species occurred across a wide range of peridomestic landscape conditions. This suggests that multiple potential vector species may be present across heterogeneous landscapes, although whether this affects *Leishmania* transmission remains to be tested with infection and host data. Land-cover composition, forest configuration, and elevation accounted for about half of the explained variance in species occurrence and abundance, but the models explained a modest share of total variation, and responses differed among species. Grassland, artificial surfaces, and forest edges were the clearest landscape correlates, which suggests that land-use decisions may shape vector assemblages and that surveillance and control strategies could benefit from integrated input from public health, agriculture, and land-use planning. Future studies that combine pathogen screening, host communities, finer-scale habitat characteristics, and short-term climatic variability would help determine whether these patterns translate into differences in vector abundance and human exposure. By combining metacommunity analyses with joint species distribution models, this study offers a community-level framework for examining how landscape heterogeneity relates to sand fly assemblages, extending beyond single-species approaches.

## Data availability

The R-script and raw dataset supporting the results of this work are available at Zenodo Digital Repository. https://doi.org/10.5281/zenodo.23366256

## Ethics approval and consent to participate

We conducted sand fly sampling in accordance with institutional and national ethical regulations. Research protocols were approved by the University of Georgia Institutional Animal Care and Use Committee (IACUC; Protocols A2023 07-015-Y2-A0 and A2022 07-008-Y3-A0), the Instituto Conmemorativo Gorgas de Estudios de la Salud IACUC (Nota 001/CIUCAL/ICGES22), and the Panamanian Ministry of Environment (MiAMBIENTE; Permit ARBG-009-2022).

## Consent for publication

Not applicable

## Declaration of interests

The authors declare no competing interests.

## Author contributions

Conceptualization: Nicole L. Gottdenker, Juliana Hoyos, Chystrie Rigg, Luis Fernando Chaves, José Calzada.

Data curation: Chystrie Rigg

Formal analysis: Juliana Hoyos, Chystrie Rigg

Funding acquisition: Nicole L. Gottdenker, Luis Fernando Chaves, Jose Calzada, Azael Saldaña, Susan Tanner, Julie Velasquez Runk, Sonia Altizer, J. P. Schmidt

Methodology: Chystrie Rigg, Nicole L. Gottdenker, Luis Fernando Chaves, José E. Calzada, Azael Saldaña, Juliana Hoyos, Vanessa Vasquez, Milixa Perea

Supervision: Nicole L. Gottdenker, Sonia Altizer, Jose Calzada Visualization: Juliana Hoyos, Nicole L. Gottdenker.

Validation: Juliana Hoyos

Writing – original draft: Juliana Hoyos, Chystrie Rigg, Nicole Gottdenker.

Writing – review & editing: Juliana Hoyos, Chystrie Rigg, Nicole L. Gottdenker, Luis Fernando Chaves, Vanessa Vasquez, Milixa Perea, Jose E. Calzada, Azael Saldaña, Vanessa Pineda, Kadir Gonzalez, Sonia Altizer, Raissa Nogueira de Brito, Adelys Reina, J. P. Schmidt, Susan Tanner, Julie Velásquez Runk.

## Supporting information

**S1 Fig. Correlogram between landscape covariates calculated at 300 m buffers around CDC Traps**. Values in cells show Pearson correlation coefficients.

**S2 Fig. Graphical summary of the JSDM statistical framework adjusted to our study on Sand fly communities in Panama.** Bullet points and green boxes correspond to data. Blues ellipses to estimated parameters.

S3 Fig. Rarefaction curves of species richness among the 12 communities visited between 2022 and 2023. (Blue horizontal lines correspond to Chao1 estimates for each community).

**S4 Fig. Box plots illustrating Shannon index (alpha diversity indices) per Communities during Dry and wet seasons**. Median values and inter-quartile ranges have been indicated in the plot.

**S5 Fig. Convergence diagnostics for the JSDM for abundance using Gelman–Rubin R ^.** Histogram shows the distribution of univariate potential scale reduction factors. Overall, most parameters fell at or below the 1.1 threshold, supporting satisfactory MCMC convergence across blocks.

**S6 Fig. Convergence diagnostics for the JSDM for occurrence using Gelman–Rubin R ^.** Histogram shows the distribution of univariate potential scale reduction factors. Overall, most parameters fell at or below the 1.1 threshold, supporting satisfactory MCMC convergence across blocks.

**S7 Fig. Effective sample size (ESS) diagnostics for the JSDM for abundance.** Overlaid histograms show the distribution of ESS values. Vertical reference lines mark common heuristics for adequate mixing (100 and 200, dashed) and a more conservative target (400, dash-dot).

**S8 Fig. Effective sample size (ESS) diagnostics for the JSDM for occurrence.** Overlaid histograms show the distribution of ESS values. Vertical reference lines mark common heuristics for adequate mixing (100 and 200, dashed) and a more conservative target (400, dash-dot).

**S9 Fig. Proportion of conditional variance explained for the JSDM abundance model.** Each bar represents the variance explained for an individual sand fly species, arranged in decreasing order of total explained variance (bar length). Anthropportunists species are marked with asterisks.

**S10 Fig. Proportion of total variance explained (Tjur R2) for the JSDM occurrence model.** Each bar represents the variance explained for an individual sand fly species, arranged in increasing order of total explained variance (bar length). Anthropportunists species are marked with asterisks.

**S11 Fig. Comparison of model predictive performance, based on estimated AUC per sand fly species.** Anthropportunists species are marked with asterisks.

**S12 Fig. Chord diagram depicting sand fly species co-occurrence based on residual associations.** The connections (arcs) between species indicate how frequently they co-occur, with thicker arcs representing more frequent co-occurrences.

**S1 Table. The variance inflation factors (VIF) of the covariates used for the Joint species Distribution Models.**

## Notes

### Competing Interest Statement

The authors have declared no competing interest.

## References

1. Johnson PT, de Roode JC, Fenton A. Why infectious disease research needs community ecology. Science. 2015;349(6252):1259504. doi:10.1126/science.1259504.

2. de Souza WM, Weaver SC. Effects of climate change and human activities on vector-borne diseases. Nat Rev Microbiol. 2024;22(8):476–491. doi:10.1038/s41579-024-01026-0.

3. Roche B, Guégan JF. Ecosystem dynamics, biological diversity and emerging infectious diseases. C R Biol. 2011;334(5-6):385–392. doi:10.1016/j.crvi.2011.02.008.

4. Chaves LF, Añez N. Nestedness patterns of sand fly (Diptera: Psychodidae) species in a neotropical semi-arid environment. Acta Trop. 2016;153:7–13. doi:10.1016/j.actatropica.2015.10.001.

5. Yamada K, Valderrama A, Gottdenker N, Cerezo L, Minakawa N, Saldaña A, et al. Macroecological patterns of American cutaneous leishmaniasis transmission across the health areas of Panamá (1980-2012). Parasite Epidemiol Control. 2016;1(2):42–55. doi:10.1016/j.parepi.2016.03.003.

6. Guo F, Bonebrake TC, Gibson L. Land-use change alters host and vector communities and may elevate disease risk. EcoHealth. 2019;16(4):647–658. doi:10.1007/s10393-018-1336-3.

7. Chaves LF. Phlebotomine sand fly species co-occurrence at the local scale: differences between agricultural and forested areas. Bol Malariol Salud Ambient. 2011;51(1):35–39. Available from: http://ve.scielo.org/scielo.php?script=sci_arttext&pid=S1690-46482011000100004&lng=es&tlng=en.

8. Cecílio P, Cordeiro-da-Silva A, Oliveira F. Sand flies: basic information on the vectors of leishmaniasis and their interactions with Leishmania parasites. Commun Biol. 2022;5:305. doi:10.1038/s42003-022-03240-z.

9. Ready PD. Biology of phlebotomine sand flies as vectors of disease agents. Annu Rev Entomol. 2013;58:227–250. doi:10.1146/annurev-ento-120811-153557.

10. Pinto Moraes JL, Marinho Santana HT, da Conceição Abreu Bandeira M, Macário Rebêlo JM. Effects of forest degradation on the sand fly communities of northeast Brazil. J Vector Ecol. 2020;45(1):89–99. doi:10.1111/jvec.12376.

11. Massey AL, Ferreira da Silva DJ, Vieira CJDSP, Allen JM, Canale GR, Bernardo CSS, et al. Using iDNA to determine impacts of Amazonian deforestation on Leishmania hosts, vectors, and their interactions. PLoS Negl Trop Dis. 2025;19(3):e0012925. doi:10.1371/journal.pntd.0012925.

12. Vivero RJ, Torres-Gutierrez C, Bejarano EE, Peña HC, Estrada LG, Florez F, et al. Study on natural breeding sites of sand flies (Diptera: Phlebotominae) in areas of Leishmania transmission in Colombia. Parasit Vectors. 2015;8:116. doi:10.1186/s13071-015-0711-y.

13. Mascari TM, Hanafi HA, Jackson RE, Ouahabi S, Ameur B, Faraj C, et al. Ecological and control techniques for sand flies (Diptera: Psychodidae) associated with rodent reservoirs of leishmaniasis. PLoS Negl Trop Dis. 2013;7(9):e2434. doi:10.1371/journal.pntd.0002434.

14. Chaves LF, Cohen JM, Pascual M, Wilson ML. Social exclusion modifies climate and deforestation impacts on a vector-borne disease. PLoS Negl Trop Dis. 2008;2(1):e176. doi:10.1371/journal.pntd.0000176.

15. Dantas-Torres F, Sales KG, Miranda DE, da Silva FJ, Figueredo LA, de Melo FL, et al. Sand fly population dynamics and cutaneous leishmaniasis among soldiers in an Atlantic Forest remnant in northeastern Brazil. PLoS Negl Trop Dis. 2017;11(2):e0005406. doi:10.1371/journal.pntd.0005406.

16. Loaiza JR, Rovira JR, Sanjur OI, Zepeda JA, Pecor JE, Foley DH, et al. Forest disturbance and vector transmitted diseases in the lowland tropical rainforest of central Panama. Trop Med Int Health. 2019;24(7):849–861. doi:10.1111/tmi.13244.

17. Ministerio de Salud de la República de Panamá. Informe consolidado de las acciones de control vectorial para las enfermedades malaria, Aedes, Chagas, leishmaniasis y roedores año 2023 [Internet]. Panamá: Ministerio de Salud; [date unknown]. Available from: https://www.minsa.gob.pa/sites/default/files/publicacion-general/034-2023_actualizacion_de_actas_de_inspeccion_reemplaza_comunicado_009-2023.pdf.

18. Dutari LC, Loaiza JR. American cutaneous leishmaniasis in Panama: a historical review of entomological studies on anthropophilic Lutzomyia sand fly species. Parasit Vectors. 2014;7:218. doi:10.1186/1756-3305-7-218.

19. Valderrama A, Tavares MG, Andrade Filho JD. Anthropogenic influence on the distribution, abundance and diversity of sandfly species (Diptera: Phlebotominae: Psychodidae), vectors of cutaneous leishmaniasis in Panama. Mem Inst Oswaldo Cruz. 2011;106(8):1024–1031. doi:10.1590/S0074-02762011000800021.

20. Rigg CA, Perea M, González K, Saldaña A, Calzada JE, Gao Y, et al. Diversity, co-occurrence, and nestedness patterns of sand fly species (Diptera: Psychodidae) in two rural areas of western Panamá. Insects. 2021;12(2):113. doi:10.3390/insects12020113.

21. Nieves E, Oraá L, Rondón Y, Sánchez M, Sánchez Y, Rojas M, et al. Effect of environmental disturbance on the population of sandflies and Leishmania transmission in an endemic area of Venezuela. J Trop Med. 2014;2014:280629. doi:10.1155/2014/280629.

22. Valdivia HO, Zorrilla VO, Espada LJ, Perez JG, Razuri HR, Vera H, et al. Diversity, distribution and natural Leishmania infection of sand flies from communities along the Interoceanic Highway in the Southeastern Peruvian Amazon. PLoS Negl Trop Dis. 2021;15(2):e0009000. doi:10.1371/journal.pntd.0009000.

23. de Oliveira MDS, da Silva RCR, Silva ASGE, Rodrigues JC, Pimenta RCJS, Leonardo FS, et al. Land use and cover changes and sand fly (Diptera: Psychodidae) assemblages in an emerging focus of leishmaniasis. Med Vet Entomol. 2025;39(3):525–537. doi:10.1111/mve.12791.

24. Guimarães RCS, Martins-Campos KM, Farias ES, de Arruda VAB, Marialva EFS, Peixoto GM, et al. Land-use change reshapes sand fly communities: diversity loss and vector persistence in Amazonian landscapes. Diversity. 2026;18(6):339. doi:10.3390/d18060339.

25. Tikhonov G, Opedal ØH, Abrego N, Lehikoinen A, de Jonge MMJ, Oksanen J, et al. Joint species distribution modelling with the R-package Hmsc. Methods Ecol Evol. 2020;11(3):442–447. doi:10.1111/2041-210X.13345.

26. Shaw JJ. A review of Leishmania infections in American phlebotomine sand flies—are those that transmit leishmaniasis anthropophilic or anthropportunists? Parasite. 2025;32:57. doi:10.1051/parasite/2025039.

27. Quijada-Alarcón J, Rodríguez-Rodríguez R, González-Cancelas N, Bethancourt-Lasso G. Spatial analysis of territorial connectivity and accessibility in the Province of Coclé in Panama. Sustainability. 2023;15(15):11500. doi:10.3390/su151511500.

28. Young DG, Duncan MA. Guide to the identification and geographic distribution of Lutzomyia sandflies in Mexico, the West Indies, Central and South America (Diptera: Psychodidae). Memoirs of the American Entomological Institute. 1994;54:1–881.

29. Galati EAB. Phlebotominae (Diptera, Psychodidae): classificação, morfologia, terminologia e identificação de adultos. Apostila de Bioecologia e Identificação de Phlebotominae. Vol. 1. São Paulo: Universidade de São Paulo; 2015. p. 1–124.

30. Ministerio de Ambiente. Cobertura boscosa y uso del suelo Panamá, año 2021 (1:50 000). Panamá: MiAmbiente, ONUREDD, FAO; 2021. Available from: https://www.sinia.gob.pa/index.php/datos-abiertos-y-geoservicios.

31. Munstermann LE. Phlebotomine sand flies and moth flies (Psychodidae). In: Medical and veterinary entomology. 3rd ed. Elsevier; 2019. p. 191–211. doi:10.1016/B978-0-12-814043-7.00012-1.

32. Chao A. Nonparametric estimation of the number of classes in a population. Scand J Stat. 1984;11(4):265–270.

33. Leibold MA, Mikkelson GM. Coherence, species turnover, and boundary clumping: elements of metacommunity structure. Oikos. 2002;97(2):237–250. Available from: http://www.jstor.org/stable/3547414.

34. Presley SJ, Higgins CL, Willig MR. A comprehensive framework for the evaluation of metacommunity structure. Oikos. 2010;119:908–917. doi:10.1111/j.1600-0706.2010.18544.x.

35. Poos MS, Jackson DA. Addressing the removal of rare species in multivariate bioassessments: the impact of methodological choices. Ecol Indic. 2012;18:82–90. doi:10.1016/j.ecolind.2011.10.008.

36. Pearce J, Ferrier S. Evaluating the predictive performance of habitat models developed using logistic regression. Ecol Model. 2000;133:225–245.

37. Tjur T. Coefficients of determination in logistic regression models—a new proposal: the coefficient of discrimination. Am Stat. 2009;63:366–372.

38. Bauer AM, Guralnick RP, Whitehead SA, Barve N, Allen JM, Campbell LP. Land use predicts proportion of West Nile virus vector-competent mosquitoes. Ecosphere. 2024;15(2):e4771. doi:10.1002/ecs2.4771.

39. Cerini F, Vignoli L, Blust M, Strona G. Functional traits predict species co-occurrence patterns in a North American Odonata metacommunity. Ecosphere. 2023;14(12):e4732. doi:10.1002/ecs2.4732.

40. Shittu RA, Thomas SM, Roiz D, Ruiz S, Figuerola J, Beierkuhnlein C. Modeling the effects of species associations and abiotic parameters on the abundance of mosquito species in a Mediterranean wetland. Wetl Ecol Manag. 2024;32(3):381–395. doi:10.1007/s11273-024-09982-3.

41. Santini MS, Fernández MS, Cavia R, Salomón OD. Co-occurrence and seasonal and environmental distributions of the sandflies Lutzomyia longipalpis and Nyssomyia whitmani in the city of Puerto Iguazú, northeastern Argentina. Med Vet Entomol. 2018;32:197–205. doi:10.1111/mve.12283.

42. González E, Salvo A, Valladares G. Natural vegetation cover in the landscape and edge effects: differential responses of insect orders in a fragmented forest. Insect Sci. 2017;24(5):891–901. doi:10.1111/1744-7917.12377.

43. Kitron U. Landscape ecology and epidemiology of vector-borne diseases: tools for spatial analysis. J Med Entomol. 1998;35(4):435–445. doi:10.1093/jmedent/35.4.435.

44. Lambin EF, Tran A, Vanwambeke SO, Linard C, Soti V, et al. Pathogenic landscapes: interactions between land, people, disease vectors, and their animal hosts. Int J Health Geogr. 2010;9:54. doi:10.1186/1476-072X-9-54.

45. Ferraguti M, Magallanes S, Suarez-Rubio M, et al. Does land-use and land cover affect vector-borne diseases? A systematic review and meta-analysis. Landsc Ecol. 2023;38:2433–2451. doi:10.1007/s10980-023-01746-3.

46. Sjödin NE, Bengtsson J, Ekbom B. The influence of grazing intensity and landscape composition on the diversity and abundance of flower-visiting insects. J Appl Ecol. 2008;45:763–772. doi:10.1111/j.1365-2664.2007.01443.x.

47. Gao X, Huang Y, Zheng J, Xiao J, Wang H. Impact of meteorological and geographical factors on the distribution of leishmaniasis’s vector in mainland China. Pest Manag Sci. 2020;76:961–966. doi:10.1002/ps.5604.

48. Claborn D, Masuoka P, Morrow M, et al. Habitat analysis of North American sand flies near veterans returning from Leishmania-endemic war zones. Int J Health Geogr. 2008;7:65. doi:10.1186/1476-072X-7-65.

49. Sandoval-Ramírez CM, Hernández C, Teherán AA, Gutierrez-Marin R, Martínez-Vega RA, Morales D, et al. Complex ecological interactions across a focus of cutaneous leishmaniasis in Eastern Colombia: novel description of Leishmania species, hosts and phlebotomine fauna. R Soc Open Sci. 2020;7:200266. doi:10.1098/rsos.200266.

50. Capucci DC, Campos AM, Soares JVR, Ramos VDV, Binder C, Lima MA, et al. Ecology and natural infection of phlebotomine sand flies in different ecotopes and environments in the municipality of Pains, Minas Gerais, Brazil. Acta Trop. 2023;238:106789. doi:10.1016/j.actatropica.2022.106789.

51. Senanayake SC, Liyanage P, Pathirage DRK, Siraj MFR, De Silva BGDNK, et al. Impact of climate and land use on the temporal variability of sand fly density in Sri Lanka: a 2-year longitudinal study. PLoS Negl Trop Dis. 2024;18(11):e0012675. doi:10.1371/journal.pntd.0012675.

52. Casaril AE, Monaco NZN, de Oliveira EF, et al. Spatiotemporal analysis of sandfly fauna (Diptera: Psychodidae) in an endemic area of visceral leishmaniasis at Pantanal, central South America. Parasit Vectors. 2014;7:364. doi:10.1186/1756-3305-7-364.

53. Hanski I. Habitat fragmentation and species richness. J Biogeogr. 2015;42:989–993. doi:10.1111/jbi.12478.

54. Laurance WF, Nascimento HEM, Laurance SG, Andrade A, Ewers RM, et al. Habitat fragmentation, variable edge effects, and the landscape-divergence hypothesis. PLoS One. 2007;2(10):e1017. doi:10.1371/journal.pone.0001017.

55. Ribeiro JCT, Nunes-Freitas AF, Fidalgo ECC, Uzêda MC. Forest fragmentation and impacts of intensive agriculture: responses from different tree functional groups. PLoS One. 2019;14(8):e0212725. doi:10.1371/journal.pone.0212725.

56. Chust G, Pretus JL, Ducrot D, et al. Scale dependency of insect assemblages in response to landscape pattern. Landsc Ecol. 2004;19:41–57. doi:10.1023/B:LAND.0000018368.99833.f2.

57. Perrin A, Rein F, Christe P, Pellet J. Habitat fragmentation impact on insect diversity: opposing forces at patch and landscape levels. Landsc Ecol. 2025;40(6):113. doi:10.1007/s10980-025-02133-w.

58. de Oca-Aguilar ACM, Rebollar-Téllez EA, Sosa-Bibiano EI, López-Avila KB, Torres-Castro JR, Loría-Cervera EN. Effect of land use change on the phlebotomine sand fly assemblages in an emergent focus of cutaneous leishmaniasis in Yucatan, Mexico. Acta Trop. 2022;235:106628. doi:10.1016/j.actatropica.2022.106628.

59. Chaves LF, Añez N. Species co-occurrence and feeding behavior in sand fly transmission of American cutaneous leishmaniasis in western Venezuela. Acta Trop. 2004;92(3):219–224. doi:10.1016/j.actatropica.2004.08.001.

60. Calzada JE, Saldaña A, Rigg C, Valderrama A, Romero L, et al. Changes in phlebotomine sand fly species composition following insecticide thermal fogging in a rural setting of western Panamá. PLoS One. 2013;8(1):e53289. doi:10.1371/journal.pone.0053289.

61. Cornel AJ, Lee Y, Almeida APG, et al. Mosquito community composition in South Africa and some neighboring countries. Parasit Vectors. 2018;11:331. doi:10.1186/s13071-018-2824-6.

62. Cleveland CA, Dallas TA, Vigil S, et al. Vector communities under global change may exacerbate and redistribute infectious disease risk. Parasitol Res. 2023;122:963–972. doi:10.1007/s00436-023-07799-2.

63. McCreadie JW, Bedwell CR. Patterns of co-occurrence of stream insects and an examination of a causal mechanism: ecological checkerboard or habitat checkerboard? Insect Conserv Divers. 2013;6:105–113. doi:10.1111/j.1752-4598.2012.00191.x.

64. Ovaskainen O, Tikhonov G, Norberg A, Blanchet FG, Duan L, Dunson D, et al. How to make more out of community data? A conceptual framework and its implementation as models and software. Ecol Lett. 2017;20(5):561–576. doi:10.1111/ele.12757.

65. Williams-Subiza EA, Brand C, Miserendino ML. Metacommunity structure analysis reveals nested patterns in deconstructed macroinvertebrate assemblages. Ecol Entomol. 2020;45:1284–1295. doi:10.1111/een.12912.

